# Where we perceive before we look: The distribution of presaccadic attention assessed with dynamic 1/f noise

**DOI:** 10.1101/2022.01.05.475160

**Authors:** Nina M. Hanning, Heiner Deubel

**Affiliations:** Allgemeine und Experimentelle Psychologie, Ludwig-Maximilians-Universität München, München, Germany; Department of Psychology and Center for Neural Science, New York University, New York, NY, USA; Institut für Psychologie, Humboldt Universität zu Berlin, Berlin, Germany

**Author notes:** corresponding author: Nina M. Hanning.

**Keywords:** presaccadic attention, spatial attention, saccadic eye movements, visual perception

## Abstract

Already before the onset of a saccadic eye movement, we preferentially process visual information at the upcoming eye fixation. This ‘presaccadic shift of attention’ is typically assessed via localized test items, which potentially bias the attention measurement. Here we show how presaccadic attention shapes perception from saccade origin to target when no scene-structuring items are presented. Participants made saccades into a 1/f (“pink”) noise field, in which we embedded a brief orientation signal at various locations shortly before saccade onset. Local orientation discrimination performance served as a proxy for the allocation of attention. Results demonstrate that (1) saccades are preceded by shifts of attention to their goal location even if they are directed into an unstructured visual field, but the spread of attention, compared to target-directed saccades, is broad; (2) the presaccadic attention shift is accompanied by considerable attentional costs at the presaccadic eye fixation; (3) objects markedly shape the distribution of presaccadic attention – demonstrating the relevance of an item-free approach for measuring attentional dynamics across the visual field.

## Introduction

We experience the world by making saccadic eye movements. Every few hundred milliseconds, a saccade shifts our gaze and focus of attention to a new piece of visual information. Interestingly, spatial attention reaches the future eye fixation already before the eyes start to move. This *‘presaccadic shift of attention’* to the saccade target is indicated by perceptual benefits (1–3) and selective modulations of sensory (orientation and spatial frequency) tuning (4–6). These behavioral correlates of presaccadic attention have been extensively characterized at the saccade target, and are described to be spatially specific, i.e., not spreading to neighboring locations (1,7). Here, we provide a broader picture by investigating the distribution of presaccadic attention across space – from saccade origin to the target, and beyond.

We used a novel full-field 1/f noise protocol (8) that allows to assess the spatial dynamics of visual attention across the field via orientation signals embedded at various locations in the noise background. Local discrimination accuracy serves as a proxy for attention deployment, and is –unique to this 1/f noise stimulus– largely independent of retinal eccentricity (8). This property enables the continuous assessment and direct comparison of attentional effects from central to peripheral locations. Moreover, the noise protocol, unlike conventional paradigms (7), does not rely on discrete test items, which pose the risk of shaping the spread of presaccadic attention by structuring the visual scene (9,10). Due to the paradigm’s dynamic nature (the 1/f noise field is continuously changing), test signal presentation does not interrupt saccade programming – a problem observed with conventional discrimination protocols that rely on sudden-onset stimuli (7). Using a 1/f noise protocol, we could measure the unbiased distribution of attention to evaluate the presaccadic shift and spread of attention, and how it is shaped by the presence of a saccade target structure.

## Results

In **Experiment 1**, we trained participants to perform horizontal saccades of approximately 7.5° into a 1/f noise field, without target objects being presented, and compared their presaccadic orientation discrimination performance – a proxy for the allocation of attention – to their performance during continued eye fixation (**Fig. 1A** & **Methods**). During *fixation* (**Fig. 1B**, blue), discrimination performance was highest for orientation signals occurring at the center of gaze (0°:80.24±5.45%; mean±SEM), and decreased with retinal eccentricity. In contrast, during *saccade* preparation (~100ms prior to saccade onset; **Fig. 1B**, green), performance was highest at the (unmarked) saccade goal (7.5°:77.43±4.33%). Thus, presaccadic attention shifted to the saccade goal even in the absence of an actual motor target. Interestingly, the attention shift was not limited to the 7.5° saccade goal; also at neighboring locations (4.5°-9°) performance was enhanced compared to fixation (4.5°:69.21±4.85%, *p*=0.0260; 6°:72.43±5.15%; *p*=0.0160; 7.5°:77.43±4.33%, *p*=0.0050; 9°:75.18±4.22%; *p*=0.0470). This spread is surprising, as presaccadic attention is often described to be narrowly focused on the saccade target (e.g., 1,7).

**Figure 1.**
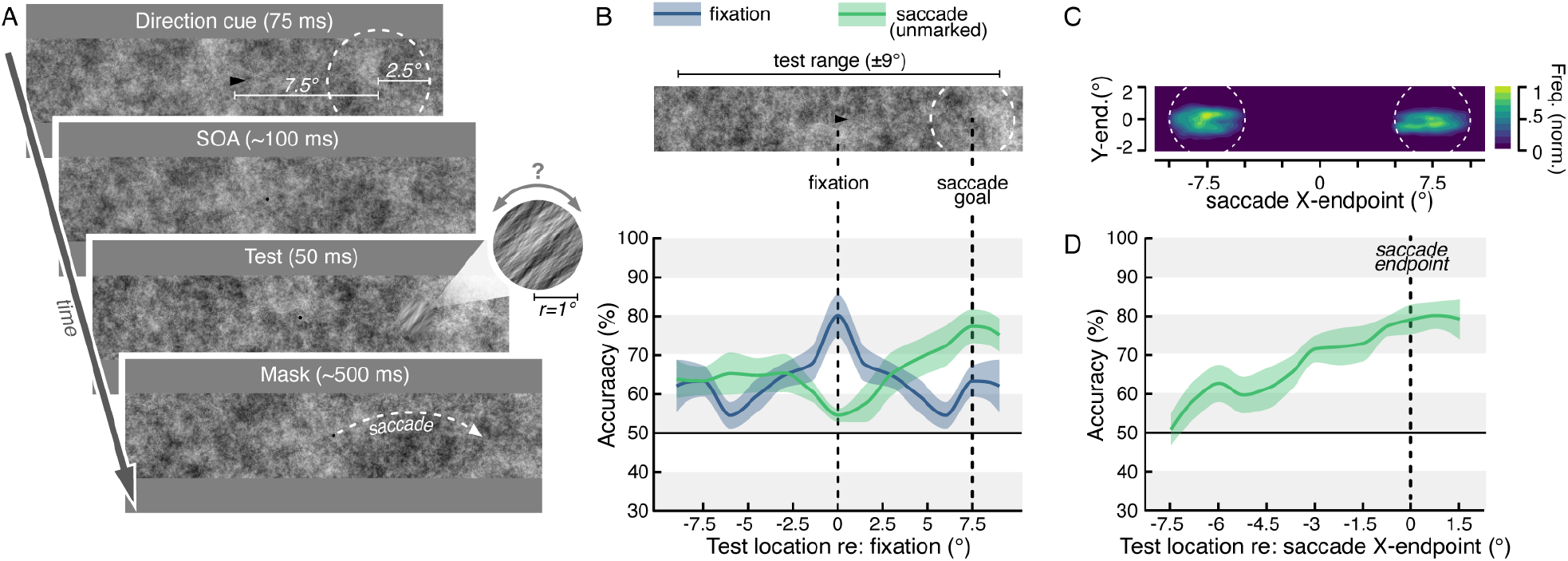
Visuospatial attention during fixation and saccade preparation. (**A**) Task. Participants fixated a central fixation dot on dynamic 1/f noise background and either made a 7.5° saccade in cue direction into the unstructured field (*saccade* trials) or kept fixating (*fixation* trials; ~15%, randomly intermixed). Shortly before saccade onset, a brief orientation signal was embedded at a random location on the horizontal (±9° around fixation). Participants indicated its tilt (here clockwise) at the end of the trial. (**B**) Group average discrimination performance relative to central fixation. Data mirrored (*fixation*) or flipped to represent rightwards saccades (*saccade*). (**C**) Normalized saccade landing frequency maps averaged across participants. (**D**) Group average discrimination performance relative to saccade endpoint. Colored areas (*B&D*) indicate ±1 SEM.

Notably, the absence of a saccade target caused considerable saccade landing variance (**Fig. 1C**). This, however, does not explain the atypically broad presaccadic attention benefit: When accounting for motor variance (**Fig. 1D**), the performance benefit remained widespread, i.e., was likewise not specially focussed on the saccade endpoint. Saccades thus draw attention to their goal even when directed into an unstructured visual field – albeit with lower spatial selectivity.

Our data moreover reveal that saccade preparation comes at a cost: Whereas discrimination performance peaked at the center of gaze during fixation, we observed the weakest performance there right before saccade onset: Foveal performance was significantly reduced from 0 to 1.5° compared to fixation (0°: 54.57±1.68%, *p*=0.0010; 1.5°:57.46±4.67%; *p*=0.0460). The presaccadic shift of attention, thus, is accompanied by a removal of processing resources from the presaccadic center of gaze.

To quantify the impact of objects on the spread of presaccadic attention, in **Experiment 2** we marked potential saccade target locations and central fixation in half of the experimental blocks (**Fig. 2A** & **Methods**). When the saccade target was *unmarked* **(Fig. 2B**, green), as in Experiment 1, presaccadic discrimination performance was highest at the saccade goal (8°:83.69±4.64%), but the benefit spread and performance was similarly enhanced at the neighboring location (6°:77.17±4.44%, *p*=0.174). In contrast, when the saccade target was *marked*, we observed a spatially far more specific presaccadic attention shift: The performance benefit was centered at the saccade target, and decreased steeply for the neighboring locations (**Fig. 2B**, purple): Peak performance at the marked target location was significantly higher (8°: 87.67±2.93%) than at the adjacent locations not overlapping with the marker (6 °:64.55±3.66%, *p* =0.002; 10°:55.84±4.34%; *p*=0.002). When directly contrasting discrimination accuracy for marked and unmarked trials from fixation to saccade goal, performance was significantly higher between 3° and 5° when no markers were presented (0.005<*p*<0.035) – further highlighting a broad, unfocussed presaccadic attention shift in the absence of target items.

**Figure 2.**
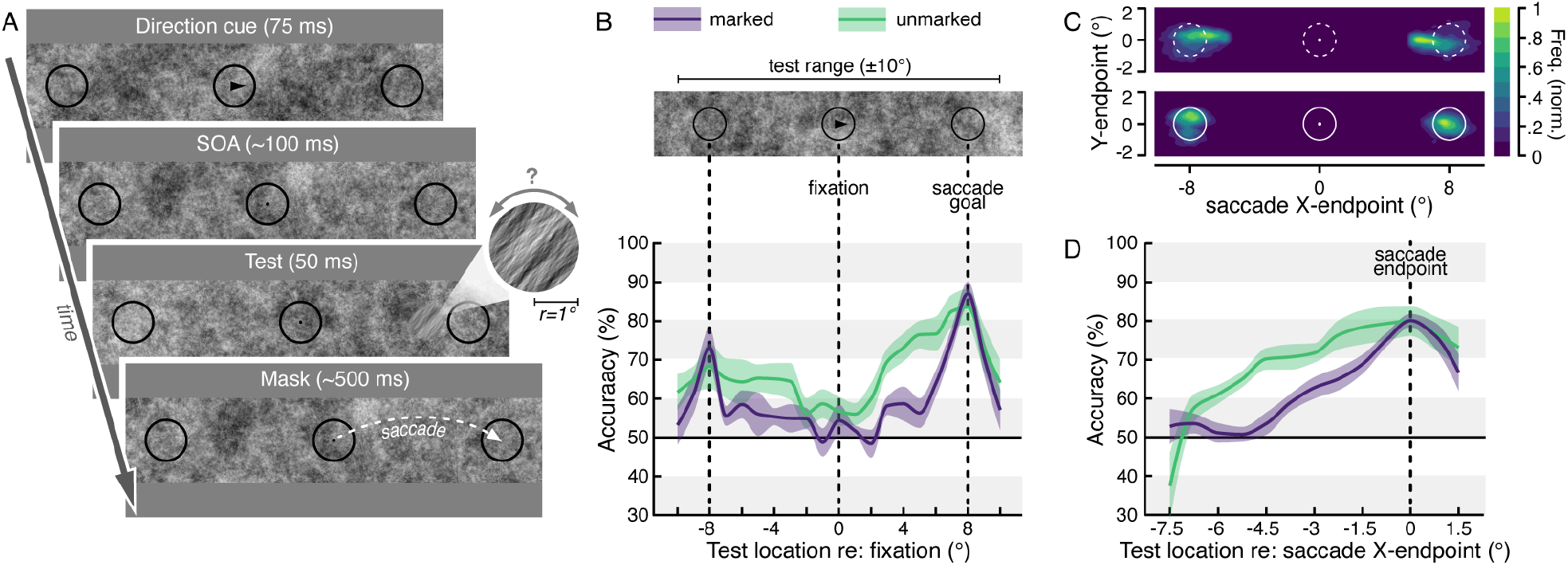
The influence of objects on presaccadic attention. (**A**) Task. Participants prepared a 8° saccade in arrow cue direction. In 50% of the experimental blocks, central fixation and the two potential target locations were framed (*marked*), in the other half no frames were presented (*unmarked*). Shortly before saccade onset, a brief orientation signal was embedded at a random location (±10° around fixation; within or between the frames). Participants discriminated its orientation after the saccade. (**B**) Group average discrimination performance relative to central fixation. Data flipped to represent rightward saccades. (**C**) Normalized saccade landing frequency maps averaged across participants for unmarked (top) and marked blocks (bottom). (**D**) Discrimination performance relative to saccade endpoint. Colored areas (*B*&*D*) indicate ±1 SEM.

Saccades made to a marked target landed closer to and scattered less around the indicated location than saccades directed to an unmarked goal (euclidean distance *marked*: 0.82±0.05° vs. *unmarked*: 1.17±0.08°, *p*=0.001; endpoint variance *marked*: 0.19±0.01° vs. *unmarked*: 0.30±0.02°, *p*=0.004; **Fig. 2C&D**). Yet again, the lower accuracy and precision of saccades in trials without marked motor target did not explain the broader spread of visual attention. When comparing performance around the endpoint of saccades made to marked and unmarked targets, the presaccadic benefit was spatially more focused when a target was presented (**Fig. 2D** purple vs. green). Accuracy for both types of trials peaked at the actual landing position (*marked* 0°:80.58±1.91%; *unmarked* 0°: 81.18±3.88%), yet when contrasting performance across space, it was significantly higher −6° to −3° relative to the endpoint when no markers were presented (0.004<*p*<0.044).

The shaping effect of markers on the spatial distribution of attention furthermore becomes evident at the −8° location, opposite the saccade target: The mere presence of a marker –even when not a saccade target– caused a weak but spatially specific performance peak (**Fig. 2D** purple; −8°:73.11±5.02%), compared to the neighboring locations (−10°:52.01±4.29%; *p*=0.002; −6°:58.94±3.55%, *p*=0.042). No corresponding benefit was observed in unmarked trials (**Fig. 2D** green; −8°:68.07±5.64%; vs. −10°:61.14±4.63%; *p*=0.282; vs. −6°: 64.87±6.12%, *p*=0.999).

## Discussion

We investigated presaccadic visual processing using a 1/f noise paradigm (8) in which local test signals are seamlessly embedded in dynamically changing background noise. This way, participants cannot anticipate potential test locations or times, and the unbiased spread of presaccadic attention can be evaluated in the absence of scene-structuring items.

Consistent with previous work, we show that during saccade preparation attention is selectively allocated towards the saccade target (1–7). Crucially, our results document that saccades are preceded by shifts of attention even when they are not directed to a localized target, but into an unstructured field. However, in contrast to the spatially highly specific deployment of attention to a physical saccade target, the presaccadic attention shift is far less focused when no target is presented – and this is not explained by the relatively higher saccade landing variance. In line with previous research (9–11), this demonstrates that it is not the saccade endpoint that determines the spread of attention, but the presence (or absence) of scene-structuring objects.

Our present and former evidence that objects mold the distribution of visual attention (8–11) is in line with an fMRI study investigating how attention modulates perception depending on the size of the ‘attention field’ (i.e., the attended area), which was manipulated using placeholder objects (12). Similar to our study, attention field size, approximated via the spread of cortical activity measured by fMRI, narrowed when placeholders marked the test location. The distribution of attentional resources across the visual field thus is shaped by scene-structuring objects, which in turn affects both behavioral and neural responses (13).

Finally, our results reveal that the attention shift to the saccade target is accompanied by a significant removal of processing resources from the presaccadic center of gaze: While visual sensitivity increased at the saccade goal during saccade preparation, perception at the current eye fixation, where performance peaks during maintained fixation, decreased so dramatically that it even became worst. This effect is reminiscent of the phenomenon of ‘inhibition of return’ (14), where processing of a previously attended location is suppressed, presumably to free processing resources for new locations of interest. Based on our results, we propose a similar mechanism for oculomotor orienting: We actively sample visual information by making saccades. Once the desired information is extracted from the current eye fixation, attentional resources are withdrawn and selectively deployed to the next relevant location – the future eye fixation.

Our observation of “presaccadic foveal blindness” as well as the spread of attention around the saccade goal could not have been investigated using conventional, item-based paradigms, since the mere presentation of local test items conflates the attention measurement (8,9). The 1/f noise paradigm thus is an ideal tool to investigate the spatiotemporal dynamics of (presaccadic) attention via its effect on visual processing.

## Materials and Methods

### Participants

Sample sizes were determined based on previous work (7,15,16). 9 participants (ages 18 - 29 years, 7 female) completed **Experiment 1**, 9 participants (ages 20 - 29 years, 5 female) completed **Experiment 2**. All participants were healthy, had normal vision, and were naive as to the purpose of the experiment (except for one author, NMH). The protocols for the study were approved by the ethical review board of the Faculty of Psychology and Education of the Ludwig-Maximilians-Universität München (approval number 13_b_2015), in accordance with German regulations and the Declaration of Helsinki. All participants gave written informed consent.

### Apparatus

Gaze position of the dominant eye was recorded using a SR Research EyeLink 1000 Desktop Mount eye tracker (Osgoode, Ontario, Canada) at a sampling rate of 1 kHz. Manual responses were recorded via a standard keyboard. The experimental software was implemented in MATLAB (MathWorks, Natick, MA), using the Psychophysics (17,18) and EyeLink toolboxes (19). Participants sat in a dimly illuminated room with their head positioned on a chin rest. Stimuli were presented at a viewing distance of 60 cm on a 21-in. gamma-linearized SONY GDM-F500R CRT screen (Tokyo, Japan) with a spatial resolution of 1,024 by 768 pixels and a vertical refresh rate of 120 Hz.

### Experimental design

**Experiment 1** (see **Fig. 1A**). Participants fixated a central black (~0 cd/m2) fixation dot (radius 0.15°) on gray background (~60 cd/m2). Once stable fixation was detected within a 1.75° from fixation for at least 200 ms, rectangular dynamic 1/f noise background (24.0° x 4.0°, mean luminance ~60 cd/ m^2^; see 8 for details) was presented and remained on the screen throughout the trail. After a random fixation period (400 - 800 ms), a black (~0 cd/m2) arrow cue (length ~0.5°, height ~0.4°) was presented centrally for 75 ms, randomly pointing either leftward or rightward (*saccade* trials). Participants were trained to perform a 7.5° saccade in arrow cue direction as fast and as precise as possible. Approximately 175 ms after cue onset (this SOA was adjusted online to the average saccade latency of each individual observer, to aim for a discrimination signal presentation in the last 100 ms before saccade onset) a local orientation signal tilted 40° clockwise or counterclockwise from vertical was embedded in the background noise (between −9° and +9° relative to fixation), windowed by a symmetrical raised cosine (radius 1.0°, sigma 0.9°). Participants were informed that this test signal would appear anywhere on the horizontal, independent of cue / saccade direction. After 50 ms the orientation signal was masked by the reappearance of non-oriented noise for 500 ms; during this interval the saccade occurred. In ~15% of trials (*fixation* trials, randomly intermixed), no arrow cue was presented, and participants were instructed to keep fixating. At the end of each trial, the dynamic 1/f background noise disappeared and participants indicated via button press in a non-speeded manner whether they had perceived a clockwise or a counterclockwise orientation. They did not receive feedback on the correctness of their response.

After an initial training, participants performed 4 experimental blocks of 160 trials each. We controlled online for broken eye fixation (outside 1.75° from central fixation before the cue onset), too short (<170 ms) or too long (>400 ms) eye movement latencies, and imprecise movements (not landing within 2.5° from saccade target center). Erroneous trials were repeated in random order at the end of each block (on average 148 ± 44 trials per observer).

To ensure a consistent level of discrimination difficulty across participants, the signal’s orientation filter strength *σ* (i.e. visibility level) used in the main experiment was titrated in a pre-test for each observer. This task was identical to the main experiment *fixation* condition. For each trial, we randomly selected the orientation filter strength (*σ* = 15 to 65) and determined the filter width corresponding to 90% discrimination accuracy by fitting cumulative Gaussian functions to the discrimination performance via maximum likelihood estimation (20).

### Experiment 2

(see **Fig. 2A**). Task and timing were identical to Experiment 1 with the following differences. In 50% of the experimental blocks (*marked* blocks), three black (~60 cd/m^2^) circular frames (radius 1.2°) marked central fixation and the potential saccade targets ±8° on the horizontal relative to fixation. In the other 50% of experimental blocks, as in Experiment 1, no frames were presented (*unmarked* blocks), and participants were trained to perform a ±8° saccade into the unstructured noise field. There were no intermixed *fixation* trials. The local orientation signal was embedded in the background noise at one out of 21 evenly spaced horizontal positions from −10° to +10° retinal eccentricity (randomly chosen independent of saccade direction; including locations within the placeholders). The signal’s orientation filter width *σ* was titrated in a pre-test as described for Experiment 1.

After an initial training, participants performed 8 experimental blocks (4 *marked* and 4 *unmarked* blocks, randomly interleaved) of 190 trials each. We controlled online for broken eye (outside 1.75° from FT before the cue onset), too short (<170 ms) or too long (>400 ms) eye movement latencies, and imprecise movements (not landing within 2.5° from saccade target center). Erroneous trials were repeated in random order at the end of each block (on average 61 ± 8 *marked* trials and 86 ± 24 *unmarked* trials per observer).

### Eye data pre-processing

For both experiments, we scanned the recorded eye-position data offline and detected saccades based on their velocity distribution (21) using a moving average over twenty subsequent eye position samples. Saccade onset and offset were detected when the velocity exceeded or fell below the median of the moving average by 3 SDs for at least 20 ms. We included trials in which no blink occurred during the trial and correct eye fixation was maintained within a 1.75° radius centered on central fixation throughout the trial (Experiment 1 *fixation* trials) or until cue onset (all eye movement trials). Moreover, we only included those eye movement trials in which the initial saccade landed within 2.5° from the required target location and did neither start before test signal offset nor later than 100 ms after signal presentation. In total we included 4,896 trials in the analysis of the behavioral results (544 ± 11 trials per participant) for Experiment 1 and 11,795 trials (1,311 ± 29 trials per participant) for Experiment 2.

### Behavioral data analysis and visualization

To visualize discrimination performance across space in Experiment 1, we interpolated between the group-averaged discrimination accuracy (% correct) for each test location, separately for the *saccade* and *fixation* condition. **Fig. 1B**: 13 evenly space bins (width 0.75°) between −9° and +9° relative to central fixation; **Fig. 1D**: 13 evenly spaced bins (width 0.75°) between −7.5° and +1.5° relative to the respective horizontal saccade landing position). Similarly, for Experiment 2 we plot the interpolated group-averaged discrimination accuracy for *marked* and *unmarked* blocks. **Fig. 2B**: 21 evenly spaced test locations between −10° and +10° relative to central fixation; **Fig. 2D**: 13 evenly spaced bins (width 0.75°) between −7.5° and +1.5° relative to the respective horizontal saccade landing position).

For all statistical comparisons we used permutation tests to determine significant performance differences between two conditions or locations. We resampled our data to create a permutation distribution by randomly rearranging the labels of the respective conditions for each observer and computed the difference in sample means for 1000 permutation resamples (iterations). We then derived p-values by locating the actually observed difference (difference between the group-averages of the two conditions) on this permutation distribution, i.e. the p-value corresponds to the proportion of the difference in sample means that fell below or above the actually observed difference (*p*-values were Bonferroni-corrected for multiple comparisons).

### Data deposition

Eyetracking and behavioral data will be available via the OSF database (https://osf.io/uzbwd) upon manuscript publication.

## Author contributions

Conceptualization and methodology: NMH, HD; Software: NMH; Investigation: NMH; Formal analysis: NMH; Visualization: NMH; Writing—original draft: NMH; Writing—review & editing: NMH, HD; Funding acquisition: NMH, HD.

## Acknowledgements

This research was supported by a grant of the Deutsche Forschungsgemeinschaft (DFG) to HD (DE336/5-1) and a Marie Skłodowska-Curie individual fellowship to NMH (898520). We thank the members of the Deubel lab as well as Efrain Hudnell for useful comments and discussions. The authors declare no conflict of interest.

